# APOE4 reduction in vascular-associated fibroblasts decreases cerebral amyloid angiopathy and associated vascular dysfunction

**DOI:** 10.64898/2026.10.02.756327

**Authors:** Fareeha Saadi, Yoonho Cho, Ilia V. Katritch, Brisa Lugo, Yue Wu, Leon CD Smyth, Megan Bosch, Kevin A Telfer, Krzysztof Hyrc, Samira Parhizkar, Xin Bao, Aishwarya Nambiar, Bernd H. Zinselmeyer, Song Hu, Jonathan Kipnis, Gwendalyn J. Randolph, Jason D. Ulrich, David M. Holtzman

**Affiliations:** Department of Neurology, Hope Center for Neurological Disorders, Washington University School of Medicine, St. Louis, MO, USA; Charles F. and Joanne Knight Alzheimer’s Disease Research Center, Washington University School of Medicine, St. Louis, MO, USA; Department of Biomedical Engineering, Danforth Campus, Washington University in St. Louis, St. Louis, MO, USA; Department of Physiology, Immunity Program, Biomedicine Discovery Institute, Monash University, Clayton, VIC, Australia; Department of Pathology and Immunology, Washington University School of Medicine, St. Louis, MO, USA; Center for Brain Immunology and Glia (BIG), Washington University School of Medicine, St. Louis, MO, USA

**Author notes:** These authors contributed equally to this work (Yoonho Cho, Ilia V. Katritch).

## Abstract

Cerebral amyloid angiopathy (CAA), characterized by cerebrovascular amyloid deposition, is strongly associated with Alzheimer’s disease, ischemic injury, and lobar hemorrhages. APOE4 is the strongest known genetic risk factor for CAA, yet the cellular sources of APOE4 that drive pathogenic vascular amyloid deposition remain poorly defined. Here, we identify vascular-associated fibroblasts (VAFs) as a key source of APOE4 that mediates CAA and cerebrovascular dysfunction. In post-mortem brain tissue from APOE4/E4 patients with confirmed CAA, APOE colocalizes with COL6⁺ VAFs around Aβ-laden vessels. Therefore, to study the contribution of APOE4 derived from VAFs in CAA, independently of parenchymal sources, we used 5xFAD mice in which the endogenous mouse Apoe coding exons are replaced with floxed human APOE4 sequences. This enables selective suppression of APOE4 expression in VAFs upon tamoxifen induction of Col1a2-CreERT^+^. APOE4 reduction in VAFs markedly reduced CAA, with the strongest effects observed in leptomeningeal vessels. Additionally, Aβ40, which preferentially associates with vascular amyloid, was significantly reduced in the vascular fraction. VAF APOE4 depletion resulted in fewer small microbleeds and an approximately 50% reduction in parenchymal border macrophages within the leptomeningeal compartment, whereas astrocytic reactivity remained unchanged. Functional assessment using in vivo multi-parametric photoacoustic microscopy revealed improved arterial dilation and blood flow in amyloid-laden leptomeningeal arteries. Together, these findings establish VAF-derived APOE4 as a compartment-specific driver of CAA and cerebrovascular dysfunction.

**One Sentence Summary:** Vascular-associated fibroblasts are a major source of pathogenic APOE4 that drives vascular dysfunction in cerebral amyloid angiopathy.

## INTRODUCTION

Apolipoprotein E (APOE) genotype strongly influences the burden and distribution of cerebrovascular amyloid, representing the strongest known genetic risk factor for cerebral amyloid angiopathy (CAA) (1, 2), an age-associated cerebrovascular disorder affecting at least 36% of individuals over 60 (1). CAA pathology is observed in approximately 60% of individuals with dementia and in nearly 90% of patients with Alzheimer’s disease (AD) (2–5). Deposition of amyloid-β (Aβ) within the vessel wall leads to basement membrane thickening, loss of vascular smooth muscle cells, impaired vasoreactivity, and progressive vascular dysfunction, ultimately culminating in cerebral hemorrhage and ischemia (5–9). As a result, CAA can independently contribute to cognitive impairment and represents a major cause of both spontaneous and anticoagulant-related intracerebral hemorrhage. In addition, CAA poses significant treatment-limiting vascular complications during anti-amyloid immunotherapy for AD (10, 11). Despite its clinical impact, disease-modifying therapies for CAA remain limited. The strong association between APOE genotype and CAA severity suggests that APOE-dependent vascular mechanisms may represent a tractable target for therapeutic intervention.

Extensive evidence indicates that APOE can directly interact with Aβ (12) and plays a central role in regulating amyloid aggregation and clearance in vivo (13, 14). APOE binds aggregated forms of Aβ and is a consistent component of parenchymal plaques and CAA (15, 16). Importantly, APOE is critical for fibrillar Aβ deposition in both brain parenchyma and in CAA, as amyloid-depositing mice on an ApoE knockout (KO) background have little to no amyloid deposition or CAA (14, 17–19). Consistent with this, studies from our lab demonstrate that targeting APOE with an anti-APOE antibody can reduce both vascular amyloid deposition and cerebrovascular dysfunction in vivo (20), as well as Aβ-associated tau seeding and spreading (21), underscoring APOE as a therapeutically actionable mediator of both parenchymal and CAA pathology. Studies have also shown that APOE isoforms differentially influence the timing and extent of brain amyloid deposition, with APOE4 accelerating the onset and increasing the burden of vascular amyloid, whereas APOE2 is relatively protective (19, 22). These genotype-dependent effects are thought to arise, in part, from differential modulation of Aβ aggregation and clearance. Beyond its role in amyloid aggregation, APOE4 also exerts both Aβ-dependent and Aβ-independent effects on the cerebrovasculature, including reductions in cerebral blood flow and increased blood-brain barrier permeability(23). The clinical significance of APOE-dependent vascular amyloid is further emphasized by anti-amyloid immunotherapy trials, in which antibodies targeting aggregated Aβ frequently induce amyloid-related imaging abnormalities (ARIA) (24). ARIA manifests as vasogenic edema (ARIA-E) or microhemorrhages (ARIA-H) (11) and occurs disproportionately in APOE4 carriers in a dose-dependent manner (25, 26), consistent with their increased burden of CAA and highlighting vascular amyloid as a key mediator of therapy-limiting toxicity (2). However, the specific vascular cell types through which APOE drives amyloid accumulation and cerebrovascular dysfunction in CAA remain poorly defined.

APOE is most abundantly expressed by astrocytes in the brain, although microglia significantly upregulate APOE expression during neurodegenerative conditions (27–29). ApoE lipoproteins from different cell types exhibit differences in lipidation, associated proteins, and glycosylation (28, 30, 31) that could affect ApoE’s ability to facilitate the development of CAA or CAA-related vascular dysfunction. In a recent study, we found that APOE4 KO in astrocytes, which form the glia limitans and are closely associated with cerebral blood vessels, was associated with decreased Aβ and amyloid plaques in the brain parenchyma but increased CAA burden. Despite this higher CAA burden, APOE4 depletion in astrocytes protected vascular integrity, improved vascular function, and reduced overall inflammatory gliosis (32). This suggests that APOE derived from different cell types may independently affect amyloid deposition in the parenchyma and blood vessels, as well as vascular function in these amyloid-laden vessels. Counterintuitively, the removal of the most abundant source of APOE in the brain, astrocytes, increased CAA. It remains unclear whether APOE expressed by cells in the cerebrovasculature affects CAA pathology differently from parenchymal sources. Single-cell analyses of the human AD brain vasculature reveal selective vulnerability and loss of vascular mural cell populations, including fibroblast-like cells, without proportional loss of vessel area, indicating degeneration within the vessel wall (33). These mural cell populations also exhibit some of the most pronounced transcriptional changes in AD post-mortem brains, with gene expression signatures consistent with vascular dysregulation and impaired blood flow. Interestingly, several vascular mural cell populations, including endothelial cells, vascular smooth muscle cells, and fibroblasts, express APOE in humans and mice (33, 34). In postmortem AD brains, fibroblasts express higher APOE levels than in healthy controls (33). This enrichment of APOE in fibroblasts, together with their close association with the vascular wall, suggests that these cells may be relevant contributors to APOE-dependent CAA pathology.

Here, we utilized a parenchymal plaque and CAA mouse model, 5xFAD (line 7031), that expresses human APOE4 and deposits more than 60% of Aβ in the form of CAA (35) to examine the contribution of APOE4 derived from vascular-associated fibroblasts (VAFs) in CAA. We assessed whether VAF-derived APOE4 exerted localized effects on vascular amyloid deposition and inflammation, as well as on parenchymal amyloid pathology. In parallel, we evaluated the impact of these changes on cerebrovascular structure and function.

## RESULTS

### APOE is expressed in COL6A1⁺ VAFs in human CAA

Although APOE4 is strongly linked to CAA, the cell types that drive vascular amyloid deposition remain poorly defined, especially those in the cerebrovascular compartment. VAFs are a relatively recently defined population in the CNS, positioned along the pial and penetrating vessels, and express APOE in both humans and mice (33, 34, 36). To determine whether VAFs express APOE in the human CAA brain, we examined postmortem cortical tissue from APOE4 carriers with neuropathologically confirmed CAA. Collagen type VI (COL6) was used to label VAFs, and Methoxy-X04 (MX04) was used to label fibrillar amyloid plaques. Low magnification overview of the cortical tissue revealed several MX04^+^ CAA-affected vessels, interspersed with COL6A1^+^ MX04+ cerebral vessels, as well as abundant parenchymal MX04^+^ plaques (Fig. 1A). High magnification images revealed that APOE immunoreactivity was enriched in MX04⁺ CAA-affected vessels. Notably, APOE immunoreactivity co-localized with COL6A1⁺ VAFs along MX04⁺ CAA-affected vessels (Fig. 1B, C). To further define the spatial relationship between APOE, COL6A1, and vascular Aβ, we generated 3D surface renderings in Imaris, which revealed that the APOE signal was associated with both COL6A1+ and MX04^+^ surfaces. Visualization of isolated surface reconstructions demonstrated that the APOE surfaces colocalized with COL6A1 and MX04 were spatially adjacent but largely non-overlapping in the CAA (Fig. 1B, C). Thus, COL6A1⁺ VAFs were enriched for APOE in human CAA.

**Fig. 1.**
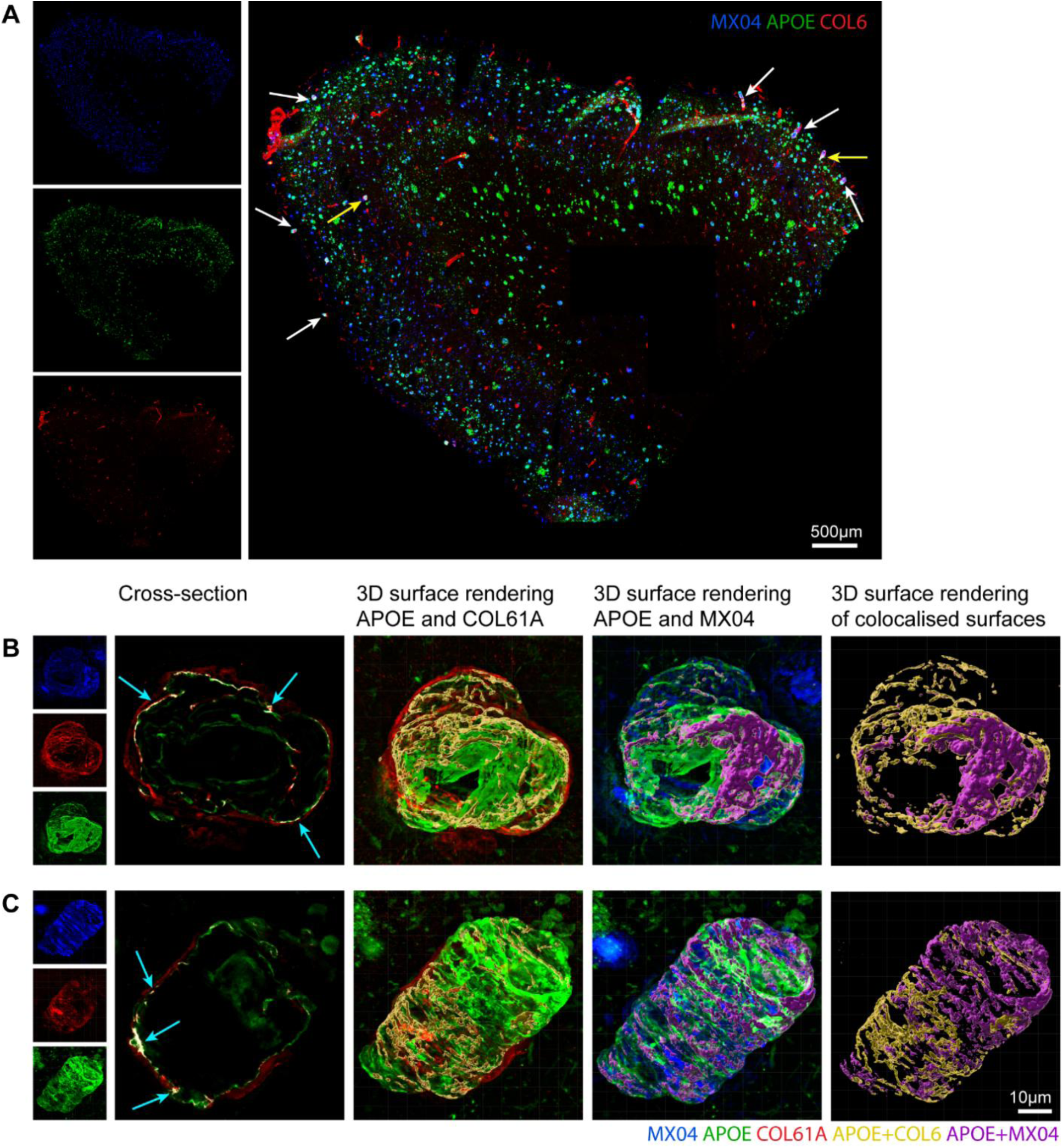
APOE colocalizes with COL6A1^+^ VAFs in human CAA patients. (A)10x overview of human parietal cortex gray matter stained for fibrillar Aβ (MX04), APOE, and VAF marker (COL6A1). White arrows indicate vessels with CAA, and yellow arrows indicate CAA vessels further magnified in B and C. (B, C) left to right, 63x magnification of vessel with CAA stained for fibrillar Aβ (MX04), APOE, and COL6A1. Cross-sectional view of a vessel depicting colocalization (white with blue arrows) of APOE and COL6A1. Representative 3D image of COL6A1 and APOE with surface rendering of colocalization (yellow). Representative 3D image of MX04 and APOE with surface rendering of colocalization (purple). Representative 3D rendering of colocalization surfaces for APOE with COL6A1 (yellow) and APOE with MX04 (purple).

### APOE4 depletion in the VAFs attenuates CAA

Our analysis of human tissue established the localization of APOE with COL6A1⁺ VAFs in CAA. Therefore, to investigate the functional relevance of APOE within this compartment, we generated 5XFAD_7031_ mice expressing human APOE4^fl/fl^ and crossed them with Col1a2-CreERT(37) mice to enable selective depletion of APOE4 in VAFs. 8-week-old, age-matched Cre^+^ and Cre^-^ mice were injected with tamoxifen intraperitoneally (i.p.) for 5 consecutive days to achieve APOE4 suppression in the VAFs (Fig. 2A). We confirmed the reduction of APOE4 expression in Slc6a13^+^ VAFs in the pial and penetrating vessels by in situ hybridization (Fig. 2B). We further performed snRNA-seq analysis on dorsal leptomeninges isolated from both groups. We found that the leptomeninges were enriched in distinct Col1a2⁺ fibroblast populations (Fig. 2C, D). Expression of canonical marker genes across clusters confirmed the annotation of distinct cell populations (S1). APOE was more abundantly expressed in pial and arachnoid fibroblast populations compared to dural fibroblasts. Both the percentage of cells expressing APOE and levels of expression showed a robust reduction upon tamoxifen treatment in the Cre⁺ group within pial and arachnoid fibroblasts (Fig. 2D), which constitute the majority of the APOE expressed by fibroblasts in the leptomeninges.

**Fig. 2.**
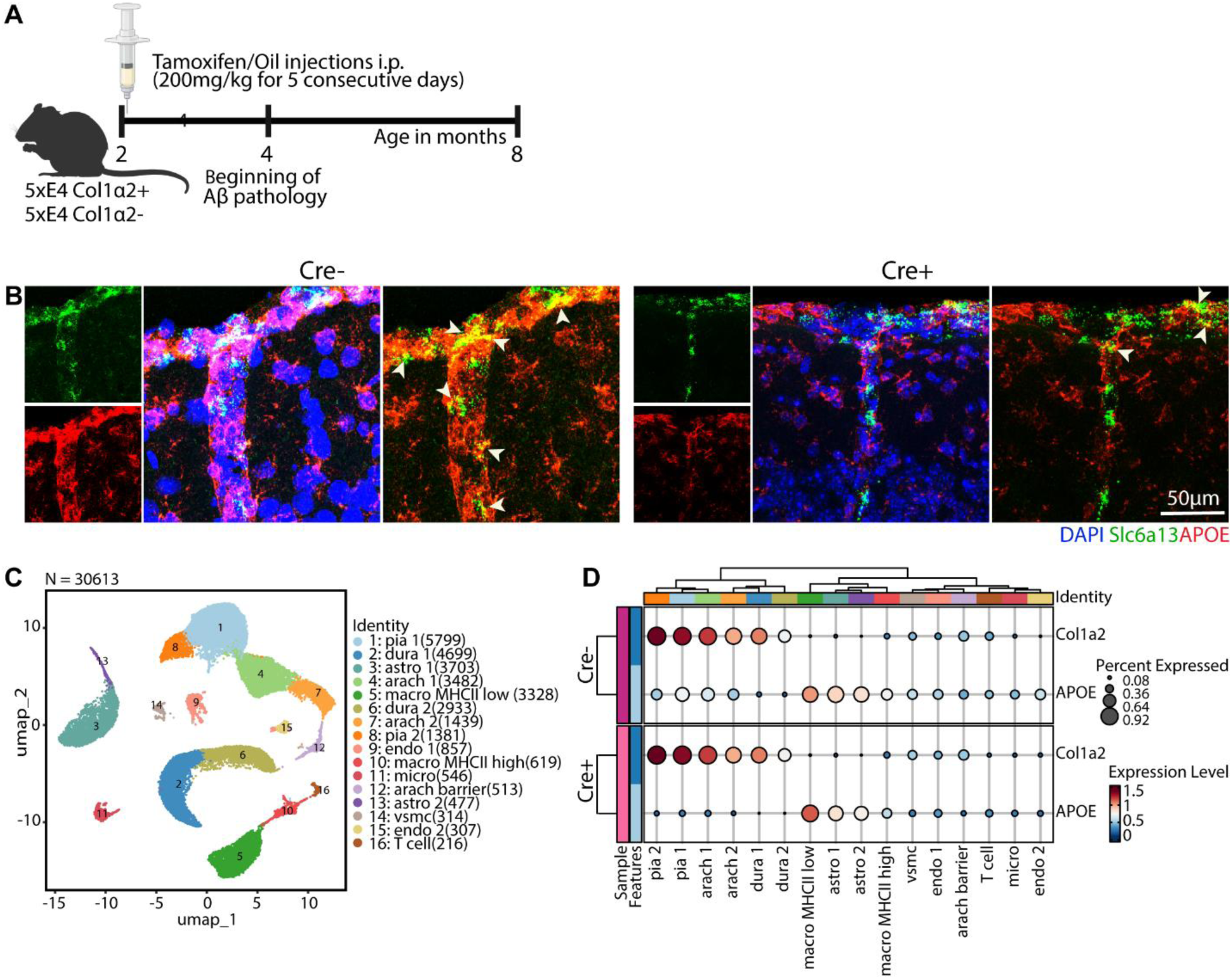
Tamoxifen-induced Cre recombination reduces APOE in the VAFs in the mouse brain. (A) Schematic of experimental design. (B) Representative ISH images from Cre⁺ and Cre⁻ mouse brains stained for VAF marker (Slc6a13), APOE, and DAPI. (C) UMAP of enriched leptomeningeal cell populations. (D) Representation of Col1a2 and APOE expression patterns across individual leptomeningeal cell populations. Key: Pia fibroblasts (Pia 1, Pia 2); Arachnoid fibroblasts (Arach 1, Arach 2); Dural fibroblasts (Dura 1, Dura 2); Astrocytes (Astro 1, Astro 2); Macrophages (MHCII low, MHCII high); Vascular smooth muscle cells (VSMC); Endothelial cells (Endo 1, Endo 2); Arachnoid barrier cells (arach barrier); T cells; Microglia (micro).

We next assessed whether local VAF-specific APOE4 depletion alters CAA burden. VAF-specific APOE4 depletion did not significantly affect total fibrillar deposits in the cortex (X34⁺) (Fig. 3A, B), although there was a significant reduction in total Aβ immunoreactivity (Fig. 3H, I). Neither fibrillar nor Aβ-immunoreactive parenchymal plaque burden was altered (Fig. 3C, J); however, we found that there was a significant reduction in total fibrillar CAA burden (Fig. 3D). Stratification by vessel compartment revealed that fibrillar CAA was reduced in both leptomeningeal (pial) and penetrating vessels (Fig. 3E, F). Consistent with this, the percentage of CAA relative to total amyloid plaque burden was significantly decreased in VAF-specific APOE4-depleted mice (Fig. 3G). Further, the assessment of Aβ immunoreactivity demonstrated a reduction in total vascular Aβ (Fig. 3K), with a strong selective decrease in leptomeningeal vascular Aβ and no detectable change in penetrating vessel Aβ (Fig. 3L, M) or the proportion of Aβ staining in the vessels (Fig. 3N). An additional Cre⁺ vehicle-treated control group exhibited comparable total amyloid burden, parenchymal plaques, and vascular amyloid relative to Cre^-^ controls upon assessment by X34 staining and Aβ immunoreactivity, relative to Cre⁻ controls, indicating that the presence of the Cre expression alone, in the absence of tamoxifen, does not alter amyloid pathology (S2).

**Fig. 3.**
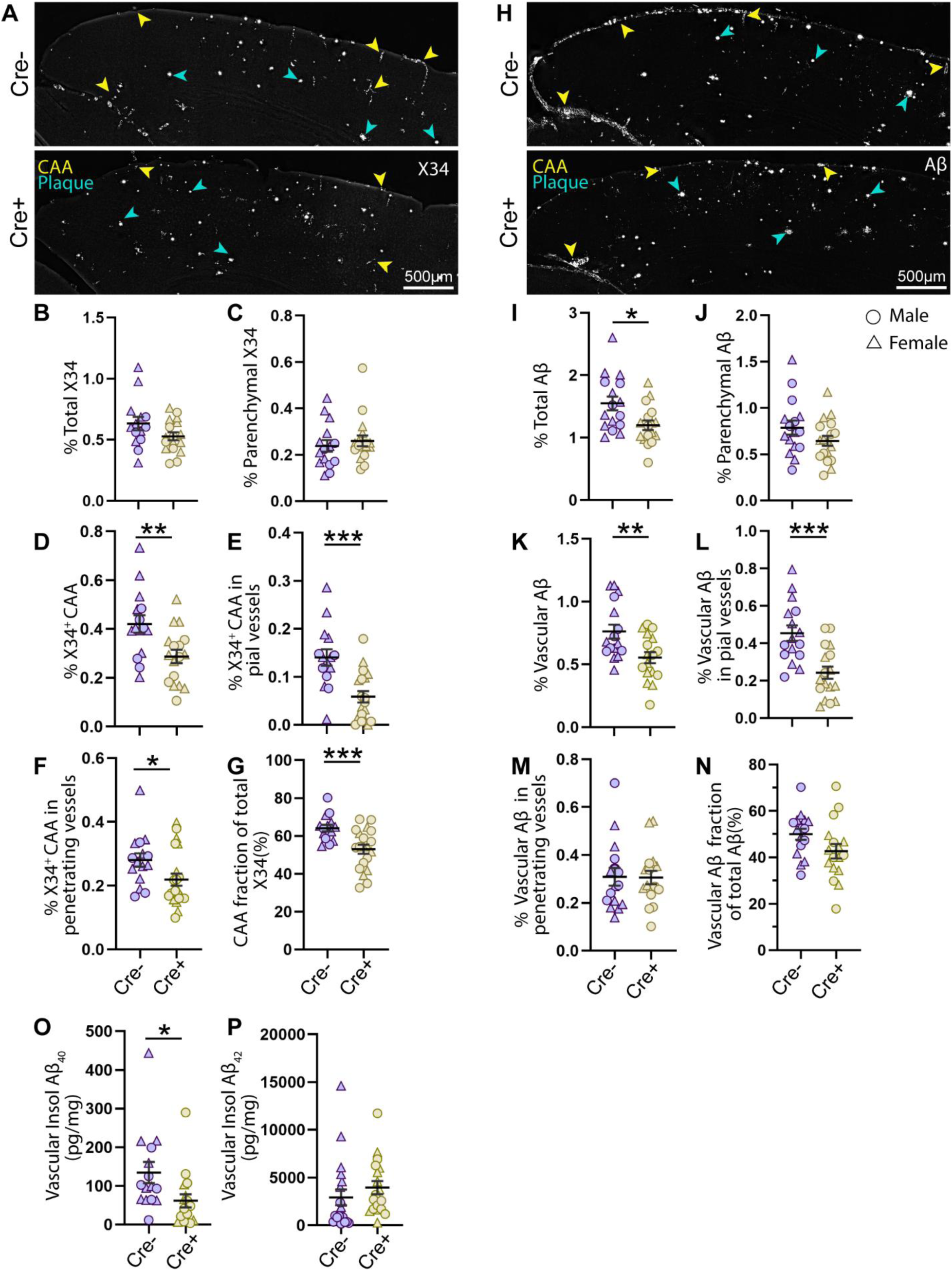
APOE4 depletion in the VAFs significantly reduces CAA. (A) Representative fluorescence images of cortices stained for fibrillar Aβ (X34). (B-G) Quantification of the percent area in the for total X34 (B), parenchymal plaques (C), total CAA (D), CAA of the pial vessels (E), CAA of the penetrating vessels (F), and proportion of CAA relative to total X34 area (G). (H) Representative images stained for Aβ immunoreactivity in the cortex. (I-N) Quantification of the percent area in the cortex for total Aβ (I), parenchymal plaques (J), total CAA (K), CAA of the pial vessels (L), CAA of the penetrating vessels (M), and proportion of CAA relative to total Aβ^+^ area (N). Yellow arrows indicate CAA, and teal arrows indicate parenchymal plaques. n=17-20 (Cre^+^) and 14-15 (Cre^-^) across analyses. (O, P) Aβ40 and Aβ42 protein levels quantified by ELISA in the formic acid soluble vascular fraction from the cortex. n = 17-18 (Cre⁺) and 15-20 (Cre⁻). n reflects the number of animals included per analysis after predefined exclusion criteria. All data expressed as means ± SEM. Student’s t-test with Welch’s correction, \**P* < 0.05; \*\**P* < 0.01; \*\*\**P* < 0.001.

Because Aβ40 is preferentially associated with vascular amyloid deposition (2, 7, 38), we quantified soluble and insoluble Aβ40 and Aβ42 in vascular and parenchymal fractions from the cortex. This analysis revealed a significant reduction in Aβ40 in the FA-soluble vascular fraction, with no change in Aβ42 in Cre⁺ mice (Fig. 3O, P). In the parenchymal fractions, no differences in either Aβ40 or Aβ42 were detected in the FA-soluble parenchymal fraction (S3A, B). Aβ40 was increased in the Hank’s Balanced Salt Solution (HBSS)-soluble fraction, whereas Aβ42 was unchanged (S3C, D) in the parenchyma. No differences in Aβ40 or Aβ42 were observed in the HBSS-soluble vascular fractions (S3E, F).

### Reduction of APOE4 in the VAFs decreases leptomeningeal parenchymal border macrophages

APOE influences immune cell recruitment and activation in the context of amyloid deposition (39–41); therefore, we next assessed how VAF-specific APOE4 depletion affects parenchymal border macrophage (PBM) and glial responses in both vascular and parenchymal compartments. We first examined PBMs (CD206^+^), which revealed a marked reduction in overall CD206⁺ area coverage in the cortex of Cre⁺ vs. Cre-mice (Fig.4A, B). No changes were observed around penetrating vessels (Fig. 4C), whereas a ∼2-fold reduction was observed in the leptomeningeal compartment (Fig. 4D). We next assessed PBM accumulation specifically around CAA. Although PBMs do not typically cluster around fibrillar plaques as microglia do, they are present in association with CAA, particularly leptomeningeal CAA (42, 43). Quantification of CD206⁺ area coverage within the perivascular region surrounding leptomeningeal CAA revealed a significant ∼50% reduction (Fig. 4E, F). We further characterized glial responses by examining microglia and astrocytic markers around CAA and parenchymal plaques. Adjacent to CAA in the penetrating vessels, an increased IBA1, TREM2, and Clec7a (Fig. 5A, B, D, E, F) immunoreactivity was observed in the Cre^+^ mice, whereas TMEM119 remained unchanged (Fig. 5A, C). Similarly, the parenchymal plaques were characterized by increased clustering of IBA1^+^ and TREM2^+^ microglia (S4 A, B, D, E), reduced TMEM119 (S4 A, C) immunoreactivity, and no significant change in Clec7a immunoreactivity (S4 D, F). In contrast, GFAP expression remained unchanged around the CAA and parenchymal plaques (S5). These results thus suggest that the selective suppression of fibroblast APOE4 modulates immune responses differently in the vascular and the parenchymal compartments.

**Fig. 4.**
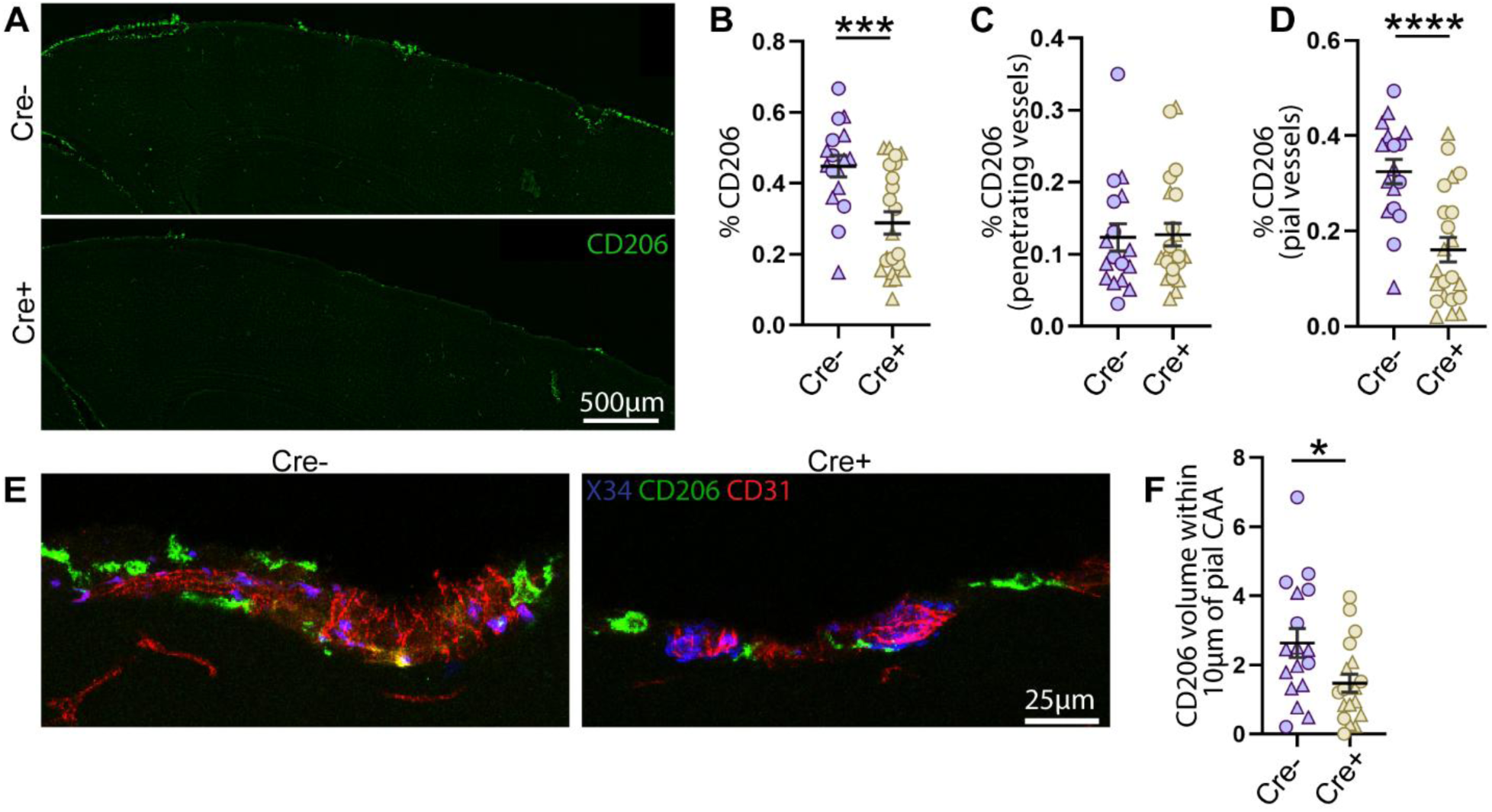
APOE4 depletion in the VAFs results in reduced PBM reactivity at the pia. (A) Representative immunofluorescence images of PBM (CD206) coverage in the cortex. (B-D) Quantification of the total % CD206 area in cortex (B), penetrating vessels (C), and pial vessels (D). n = 22 (Cre⁺) and 17 (Cre⁻). (E) Representative fluorescence image of PBM (CD206) around pial CAA. (F) Quantification of % CD206 volume within a 10μm radius of the pial CAA shown in (E). n = 19(Cre⁺) and 17 (Cre⁻). n reflects the number of animals included per analysis after predefined exclusion criteria. All data expressed as means ± SEM. Student’s t-test with Welch’s correction, \**P* < 0.05; \*\*\**P* < 0.001; \*\*\**P* < 0.0001.

**Fig. 5.**
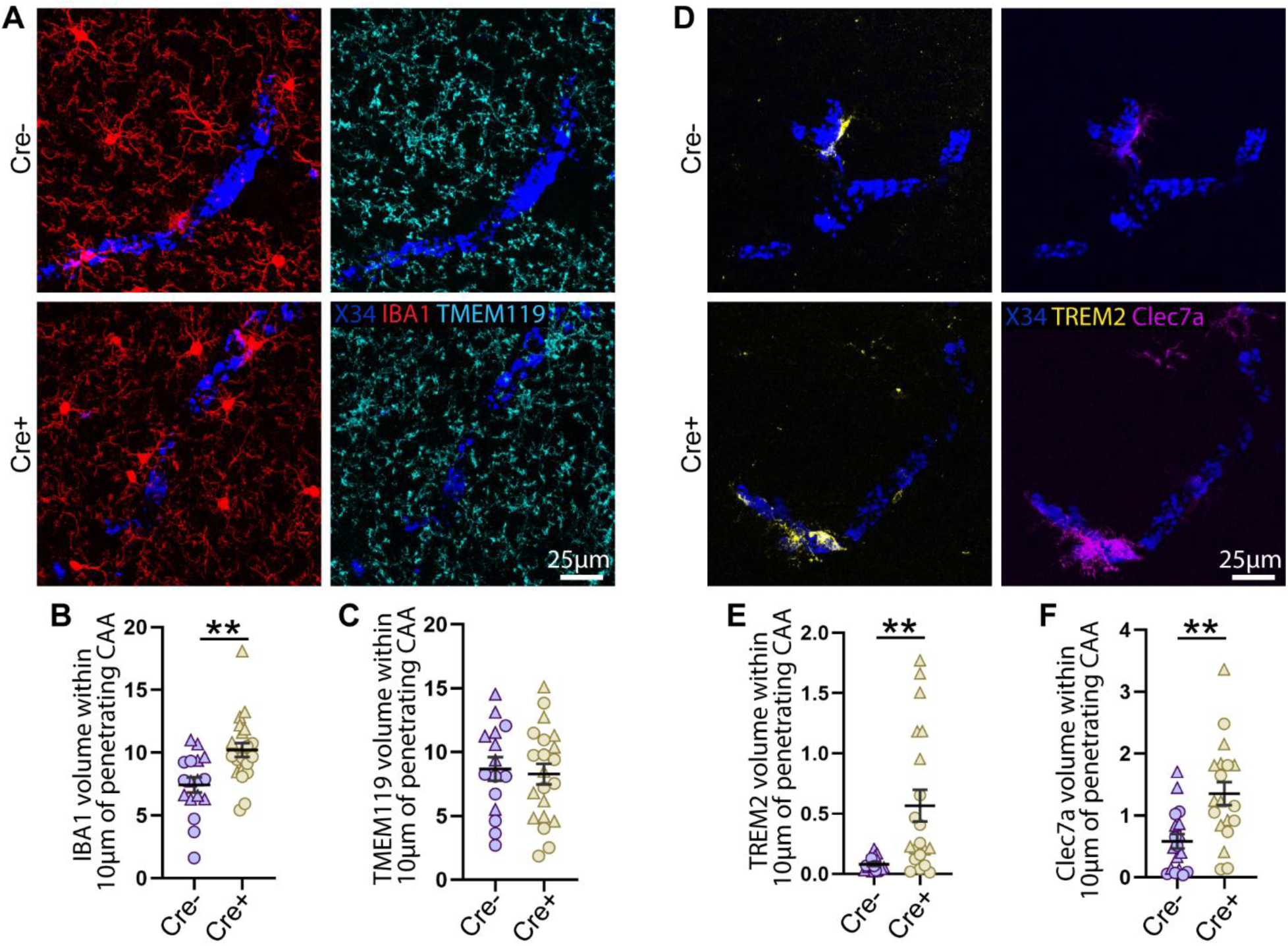
APOE4 depletion in the VAFs increases microglial reactivity around penetrating CAA. (A) Representative immunofluorescence images of fibrillar Aβ (X34) in penetrating CAA and microglial markers IBA1 and TMEM119. (B, C) Quantification of % IBA1 (B), and TMEM119 (C) volume within a 10μm radius of penetrating CAA shown in (A). n = 21-22(Cre⁺) and 15-17 (Cre⁻). (D) Representative immunofluorescence images of fibrillar Aβ (X34) in penetrating CAA and microglial markers TREM2 and Clec7a. (E, F) Quantification of % TREM2 (E), and Clec7a (F) volume within a 10μm radius of penetrating CAA shown in (D). n = 19-20(Cre⁺) and 16-18 (Cre⁻). n reflects the number of animals included per analysis after predefined exclusion criteria. All data expressed as means ± SEM. Student’s t-test with Welch’s correction, \*\**P* < 0.01.

### VAF-specific APOE4 depletion improves cerebrovascular function in leptomeningeal arteries

CAA is associated with cerebrovascular dysfunction. Because we observed a significant decline in CAA of leptomeningeal arteries upon APOE4 reduction in the VAFs, we sought to determine whether this affected vascular function. The leptomeningeal blood vessels were imaged through a cranial window using multi-parametric photoacoustic microscopy (44). Functional readouts were compared between baseline and after hypercapnia challenge (Fig. 6A). The results showed that the leptomeningeal arteries in Cre^+^ mice exhibited improved vessel dilation (Fig. 6B), increased blood flow speed (Fig. 6C), but no change in oxygen saturation in response to hypercapnia (Fig. 6D). Functional analysis of the pial veins showed improved oxygen saturation but decreased dilation and no alteration in blood flow speed in the Cre^+^ mice (S6). X34 coverage was similar between groups (median 22% vs 19%) and did not show significant correlation with vessel dilation, blood-flow speed, and oxygen saturation when photoacoustic imaging was co-registered with X34-stained widefield images, as illustrated in S7. Therefore, vascular responses were compared without stratification or normalization to X34. Microhemorrhage analysis post-mortem revealed a significant decrease in very small bleeds (<50μm) (Fig. 6E) in the Cre^+^ mice.

**Fig. 6.**
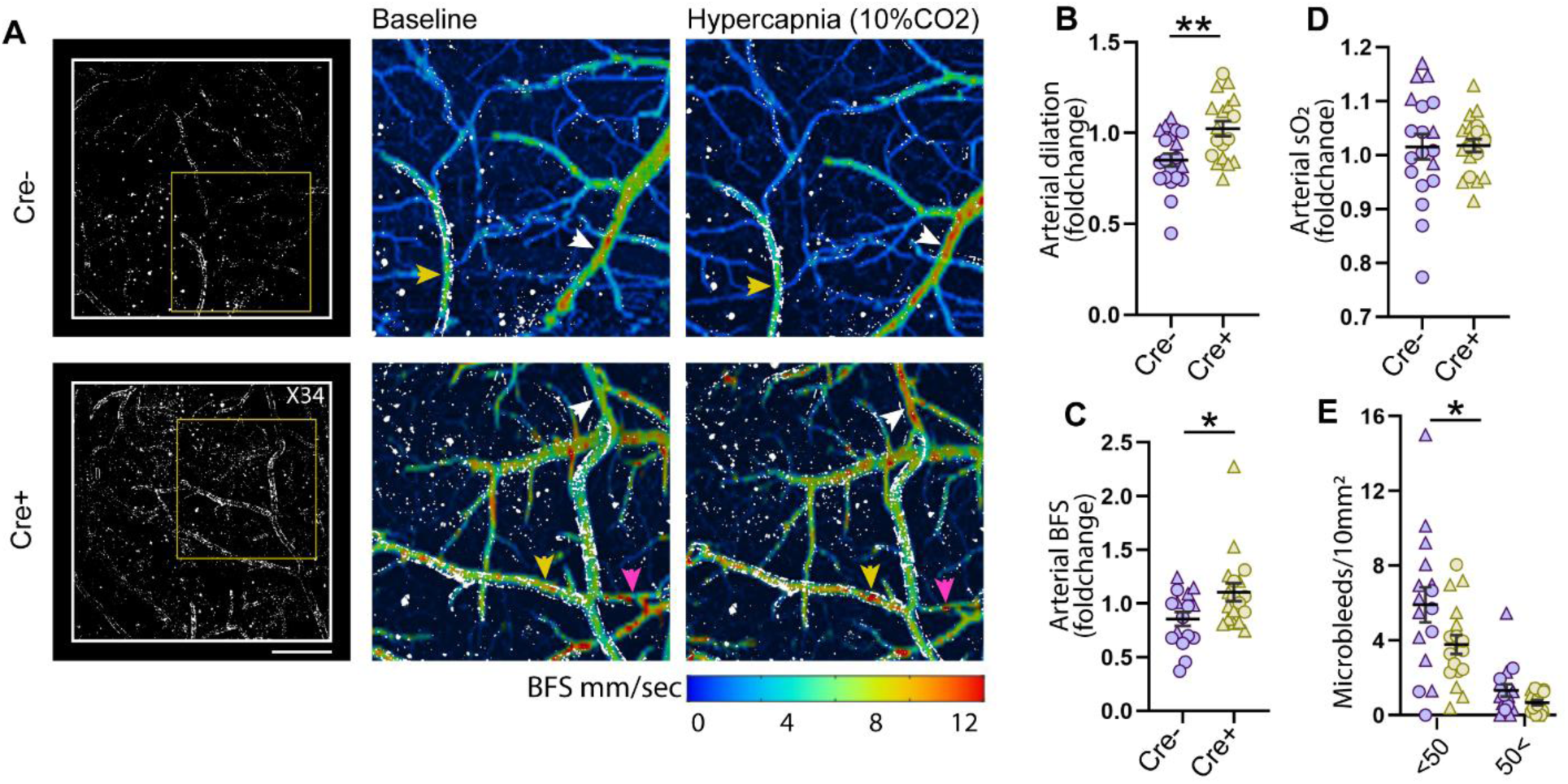
APOE4 depletion in the VAFs improves vascular function and reduces microbleeds. (A) Representative images (left to right) of pial vessels stained for fibrillar Aβ (X34), the yellow box indicates the region shown at higher magnification in the adjacent panels displaying blood-flow speed(BFS) maps at baseline and after hypercapnia obtained by multi-parametric photoacoustic microscopy, colored by flow speed with X34 overlay, scale bar, 500μm. (B-D) Quantification of fold-change in arterial dilation (B), arterial BFS (C), and arterial oxygen saturation (sO_2_) (D) upon hypercapnia relative to baseline in CAA^+^ arteries. n = 19-20 vessels from 7 mice (Cre^+^) and 16-18 vessels from 8 mice (Cre^-^). Arrows of the same color represent the same vessel at baseline and after hypercapnia. (E) Quantification of Prussian blue staining in the cortex representing microbleed frequency. n=16-20 mice in each group. n reflects the number of animals included per analysis after predefined exclusion criteria. All data expressed as means ± SEM. Student’s t-test with Welch’s correction (B–D); ordinary one-way ANOVA with Sidak’s multiple-comparison test (E). \**P* < 0.05; \*\**P* < 0.01.

## DISCUSSION

APOE is the strongest common genetic determinant of both AD and CAA(2, 7). In particular, the ε4 isoform markedly increases risk and heightens cerebrovascular vulnerability (4, 6). Although APOE is established as a central determinant of CAA pathogenesis, the cellular sources and mechanisms by which APOE4 drives vascular amyloid deposition and cerebrovascular dysfunction remain poorly defined. In mouse models that develop CAA, a global KO of APOE completely blocks CAA formation (18). We previously showed that APOE4 reduction in astrocytes was associated with markedly reduced parenchymal Aβ plaques but increased CAA burden. Astrocytes are the most abundant producers of APOE in the CNS (32); however, the predominance of astrocytic APOE does not fully explain the spatial specificity of CAA, suggesting that non-glial, vascular-adjacent cell populations may provide a locally pathogenic source of APOE4 within the vessel wall. Here, we investigated whether VAFs, a fibroblast-lineage population enriched along leptomeningeal and penetrating vessels, act as a compartment-specific source of APOE4 that locally promotes CAA.

Our findings indicate that VAF-derived APOE4 regulates vascular amyloid deposition in a highly compartment-specific manner, selectively influencing CAA, particularly within leptomeningeal vessels, without altering parenchymal plaque burden. This is consistent with enrichment of VAFs in the leptomeninges(36) and the most robust decrease in APOE in pial and arachnoid fibroblasts using the Col1a2-CreERT promoter. Although local effects of APOE on parenchymal Aβ aggregation have been described, a direct role for locally produced APOE in driving vascular amyloid deposition has not been established. Our results indicate that APOE availability within distinct vascular niches governs the CAA pathology locally, rather than globally regulating overall cerebral amyloid deposition. Parenchymal plaques predominantly deposit Aβ42, whereas CAA comprises both Aβ40 and Aβ42 (45, 46). Given the established link between CAA progression and Aβ40 accumulation, we examined whether the decrease in CAA correlated with Aβ40 levels. Indeed, biochemical analyses revealed reduced insoluble vascular Aβ40, with no corresponding changes in Aβ42 or insoluble parenchymal Aβ40/42.

This raised the question of whether the brain is clearing amyloid more effectively or whether Aβ is simply being redistributed between compartments. The concomitant increase in soluble parenchymal Aβ40 favors the latter interpretation and is most consistent with the redistribution of Aβ40 away from the vasculature. Recent studies have shown that initial Aβ seeding in parenchymal plaques requires microglia (47). However, the absence of microglia in amyloid-depositing models, while markedly decreasing parenchymal plaques, increases CAA (48). As microglia are not required for CAA formation, it may be that the myeloid population around vessels, PBMs, is either involved in Aβ seeding or clearance.

One of the most prominent results of our study was the effect of VAF-specific APOE4 depletion on PBMs along leptomeningeal vessels. We observed a ∼50% decrease in area covered by CD206^+^ PBMs specifically in this compartment, with no evident change around penetrating vessels. This reduction was localized in the CAA of the leptomeninges. Perivascular macrophages, including PBM, are increasingly implicated in the pathogenesis of CNS diseases, including CAA and AD (42, 49, 50). In CAA, they respond to vascular amyloid deposition and perivascular remodeling while also contributing to vascular stress when chronically activated (43, 51). Our findings complement this evidence by showing that PBMs respond strongly to changes in vascular amyloid burden (43, 52). Even more interestingly, astrocytic reactivity, as assessed by GFAP staining, was unaltered around the CAA, and microglial changes, though apparent, were confined to plaques and penetrating vessels, consistent with their local engagement with these pathological features. Together, these results indicate that PBMs are likely the primary immune cells that respond dynamically to changes in pial vessel CAA, reflecting their position in the perivascular space and their proximity to this pathology. However, future studies will be required to define the mechanisms governing the apparent reduction in leptomeningeal PBMs, to assess whether this is due to altered recruitment or survival, altered state/reactivity, and to determine whether PBMs are drivers of, or responders to, changes in vascular amyloid burden.

We observed clear improvements in vascular function upon APOE4 depletion from VAFs. The functional gains in the amyloid-laden pial arteries were significant, including improvements in blood vessel dilation and cerebral blood flow, and followed the reduction observed in CAA. The dissociation between vascular amyloid burden and hemodynamic measures indicates that cerebrovascular dysfunction is not strictly determined by changes in Aβ levels. Prior studies and our own work manipulating astrocytic APOE demonstrate that vascular function can improve independently of CAA burden (23, 32). These findings identify APOE4 within fibroblast-associated vascular niches as directly contributing to cerebrovascular dysfunction, and that alleviation of APOE-associated local vascular toxicity underlies functional recovery in Aβ-laden leptomeningeal vessels. Extending our findings to human disease, APOE localization to COL6A1⁺ VAFs in Aβ-laden vessels represents the first histological demonstration of APOE enrichment in this vascular compartment in the human CAA brain, thus establishing fibroblast-associated vascular niches as a conserved and functionally relevant APOE-enriched compartment in a human cerebrovascular disease.

In conclusion, our study identifies VAFs as a critical source of pathogenic APOE4, which locally regulates leptomeningeal CAA, perivascular immune responses, and cerebrovascular function. This work thus provides evidence that APOE4 targeting can be spatially constrained without inducing global inflammation or altering parenchymal amyloid burden. The spatial precision is particularly relevant in light of vascular adverse events, including ARIA, which are associated with anti-Aβ immunotherapies (1, 11, 24). Our data suggest that restricting APOE modulation to the vascular-adjacent mural cell compartments may mitigate vascular toxicity while preserving essential APOE functions in other brain cell types, potentially improving both therapeutic safety and efficacy. However, perivascular flow has been implicated in solute transport pathways involved in CAA progression and is sensitive to vessel wall compliance and basement membrane composition and function, which could be affected by modulating APOE4 in VAFs but were not assessed in this study (53–55). Another limitation of this study is that the findings from our amyloid-focused CAA model cannot be fully extrapolated to AD-associated CAA, which occurs in the context of tau-mediated neurodegeneration and cognitive decline. We have shown that an antibody against non-lipidated human APOE3 and APOE4, HAE4, can reduce parenchymal Aβ plaques, CAA, alleviate vascular dysfunction, and strongly reduce Aβ-mediated tau seeding and neuritic dystrophy(20, 21). However, extending these findings will require VAF-specific pharmacological targeting tested in models that incorporate both amyloid and tau pathology. Data from the human CAA patient brains are promising, and future studies integrating human validation, longitudinal vascular imaging, and clinically tractable targeting approaches will be essential to determine whether VAF-APOE modulation can be leveraged to reduce CAA burden and vascular risk in AD.

## MATERIALS AND METHODS

### Study design

The goal of this study was to examine whether VAF-derived APOE4 contributes to CAA formation and arteriolar dysfunction. To investigate this question, we used the 5XFAD_7031_ mice expressing human APOE4^fl/fl^, which deposit approximately 60% of Aβ in CAA and selectively suppress APOE4 expression in VAF following tamoxifen induction at 2 months of age. Littermates were genotyped and randomly assigned to experimental groups, with age and sex balanced. Animals were co-housed, and treatments were conducted by two investigators. Experiments and analyses were performed by independent investigators blinded to genotype and treatment. Sample sizes (N = 7-10 per sex per group) were determined based on prior studies from our laboratory and are sufficient to achieve ∼80% power to detect a 25% difference in Aβ load, including CAA, at p = 0.05 using a two-sided two-sample t-test assuming equal variance(32). These mice were used to assess the effects of VAF-derived APOE4 using histological analyses in the left hemisphere, biochemical measurements by ELISA in sub-dissected right-hemisphere cortical tissue, vascular function by live imaging, and snRNA-seq analysis of leptomeningeal tissue. Injections were administered in multiple batches due to the complexity of the genotypes; each batch included at least 5 mice per group, with balanced representation across all experimental groups. Data from all batches were pooled for histological and biochemical analysis, as no batch effects were observed. Samples from the same batch were used to assess vascular function and snRNA-seq of the leptomeninges. Potential outliers were identified and removed using ROUT analysis (*Q* = 1%) on GraphPad Prism. Each animal is presented as an individual biological replica, averaged from technical replicates. Details on replication and sample sizes are mentioned in the figure legends.

### Human tissue immunohistochemistry

Human parietal cortex tissue samples were acquired from the Banner Institute of Health. The sample was from an APOE4 carrier with a confirmed AD diagnosis, a Braak score of VI, and CAA. For immunofluorescent staining of APOE expression in COL6A1^+^ VAFs in human CAA, free-floating 40um brain tissues were washed with PBS^+^10U/ml heparin (PBSH) three times (10min/cycle). Following washing, tissues were delipidated in 200 mM lithium hydroxide, 10 mM lithium hydroxide, and 40 mM boric acid (pH 9) at 60 degrees Celsius for 72 hours. Sections were then washed again in PBSH (three times, 10min/cycle). Autofluorescence was quenched by incubating sections in 30% hydrogen peroxide for 1 hour at room temperature, followed by another PBSH wash (three times,10min/cycle). Antigen retrieval was performed by incubating sections in preheated Tris-EDTA buffer (173 mM Tris + 1 mM EDTA + 0.05% Tween 20) at 95 °C, then cooling to room temperature for 30min. Sections were equilibrated with three 20min PBSH washes. Tissue sections were then blocked for 1 hour at room temperature in PBS containing 10% normal donkey serum, 10% bovine serum albumin (BSA), and 0.25% Triton X-100. This was followed by a 3-day incubation at 4°C with the following primary antibodies: rabbit anti-COL6A11a (Atlas Antibodies, HPA019142; 1:100), mouse anti-APOE (Cell Signaling, E8C2U; 1:100), and methoxy-X04 (8 μM). After primary incubation, sections were washed in PBSH (three times, 10min/cycle) and incubated for 2 hours at room temperature with fluorescent-labeled secondary antibodies: donkey anti-rabbit 647, 1:200, and donkey anti-mouse 488, 1:200 (Jackson Immunoresearch). Sections were then washed in PBSH and mounted. Once air-dried, autofluorescence was quenched a second time using TrueBlack lipofuscin autofluorescence quencher (Biotium #23014) for 2 min and washed. Coverslips were applied with ProLong Gold Antifade Mounting Media (Invitrogen, P36965). Images were acquired on a Leica Stellaris confocal microscope equipped with a full-spectrum white-light laser and analyzed with Imaris (Oxford Instruments).

### Animals

All mouse experiments were approved by the Institutional Animal Care and Use Committee of Washington University in St. Louis. The 5XFAD_7031_ (56) mice expressing 2 copies of human APOE4^fl/fl^, are a model that shows Aβ pathology in the parenchyma (∼40%) and vascular Aβ deposits (∼60%) in the form of CAA. Col1a2-cre/ERT (JAX, 029235) were first crossed with APP and Presenilin-negative, APOE4flox/flox to establish Col1a2-cre/ERT APOE4^fl/fl^ breeders. Experimental 5XFAD-Col1a2-cre/ERT APOE4^fl/fl^ mice were generated by crossing 5XFAD/APOE4^fl/fl^ males with Col1a2-cre/ERT APOE4^fl/fl^ females. We only use male 5XFAD/APOE4^fl/fl^ for breeding because maternal inheritance of the transgene in the 7031 line causes significant suppression of CAA and parenchymal plaque formation(57). Littermates were assigned to tamoxifen-treated Cre-positive (Cre^+^) or Cre-negative (Cre^-^) groups, with an additional Cre^+^ oil-treated control included for selected histological analyses. Mice were maintained in a controlled environment (22 °C; 33–39% humidity) on a 12 h light–dark cycle, with ad libitum access to food and water, and no more than five animals per cage.

### Tamoxifen-induced Cre recombination

Tamoxifen was prepared at a concentration of 20 mg/mL in corn oil by shaking at 37 °C overnight, then stored at 4 °C for up to 1 month. Experimental mice received intraperitoneal injections of tamoxifen (200 mg/kg body weight) or vehicle beginning at 2 months of age for five consecutive days.

### Tissue collection

All mice were sacrificed at 8 months of age when both parenchymal plaques and CAA had begun to plateau. Mice were anesthetized with i.p. pentobarbital (100 mg/kg), followed by transcardial perfusion with ice-cold PBS containing 0.3% heparin. The skull cap was carefully removed to collect the dura; right hemibrain was sub-dissected to collect the cortex, snap frozen, and stored at −80 °C for biochemical assessments; left hemibrain was post-fixed in 4% paraformaldehyde for 48 hours, followed by cryoprotection in 30% sucrose until completely submerged. Tissues were then sectioned sagittally at 30 μm on a microtome for histological analysis and stored at −20 °C in a cryoprotectant solution (0.2 M PBS, 15% sucrose, 33% ethylene glycol).

### In situ hybridization (ISH)

30μm sagittal brain sections were mounted on charged glass slides and dried overnight at −20 °C. Multiplexed fluorescence in situ hybridization was performed the following day using the RNAscope Multiplex v2 assay kit (323100, ACDBio) according to the manufacturer’s instructions with minor modifications. Sections were dehydrated through a graded ethanol series, treated with hydrogen peroxide for 10 min at room temperature, subjected to target retrieval in boiling 1× RNAscope Target Retrieval Reagent, and incubated with Protease III for 30 min at 40 °C. Hybridization was performed using Mus musculus Slc6a13(461011, ACDBio) and human APOE (433091-C2, ACDBio) probes for 2 hours at 40 °C, followed by sequential amplification with AMP for 30 minutes each at 40 °C. The samples were then washed and coverslipped with DAPI.

### snRNA-seq analysis of leptomeninges

After perfusion, dorsal leptomeningeal tissue was dissected under a stereomicroscope and immediately placed in ice-cold RPMI. The tissue was transferred to nuclei isolation medium (NIM; 250 mM sucrose, 25 mM KCl, 5 mM MgCl₂, 10 mM Tris-HCl, Sigma-Aldrich; supplemented with protease inhibitors (Thermo Fisher Scientific), RNase inhibitor (1 U/ml, Takara), DTT (1mM, Sigma-Aldrich), and Triton X-100(0.1%v/v, Sigma-Aldrich), pH 7.5), and mechanically disrupted by fine chopping for 4-5 min. The suspension was diluted with NIM to a volume of 1ml and further homogenized by sequential douncing with loose and tight pestles, ten times each. The homogenate was filtered through a 30 µm mesh with a glass pestle, followed by repeated washes to maximize nuclei recovery. The homogenate was transferred to precoated lo-bind tubes (Eppendorf) and centrifuged at 500g for 10 min. After discarding the supernatant, the nuclei were resuspended in 5% BSA buffer supplemented with RNase inhibitor (1U/μL) and DAPI (0.2 μg /mL) and isolated by fluorescence-activated sorting based on DAPI signal using a FACS Aria II. Sorted nuclei were collected in lo-bind tubes precoated with the BSA buffer. Single-nucleus libraries were constructed using the 10x Genomics Chromium platform and the Chromium Single Cell 3′ Library & Gel Bead Kit version 3, and libraries were sequenced on the Illumina NextSeq 500 system. Sample demultiplexing, barcode processing, and single-cell 3’ counting were performed using CellRanger (10X Genomics) using a custom pre-mRNA mouse reference (mm10) plus human *APOE*. The Seurat v5 (51) and SoupX (58) R packages were used for subsequent analysis. Ambient RNA was subtracted using SoupX and low-quality nuclei removed. Samples were merged into one dataset and processed using Single Cell Transformation (SCT) to normalize reads, identify variable genes, and scale data while regressing out mitochondrial genome reads. PCA was performed, and samples were integrated using anchor-based canonical correlation analysis (CCA) integration. Top 20 PCs were used for non-linear dimension reduction (RunUMAP) and graph-based clustering (FindNeighbors, FindClusters). Non-leptomeningeal populations (ie, neurons and oligodendrocytes) were removed, and datasets were subsequently reprocessed and integrated. The final analysis used the top 15 PCs for non-linear dimensional reduction and graph-based clustering at a resolution of 0.3. Cell populations were manually annotated based on gene expression profiles. UMAPs and expression dotplots for publication were generated using scPlotter(59).

### Immunofluorescence staining

Immunofluorescence was performed on 30μm sagittal brain sections per mouse. Briefly, sections were first washed in Tris-buffered saline (TBS), then permeabilized in 0.25% TBS–Triton X-100 (TBS-X). Then, the sections were incubated in X-34 (1:5000, SML-1954, Sigma) in a 60% TBS/40% ethanol (EtOH) solution, with the pH adjusted using 1 N NaOH (1:500), for 20 min at room temperature. Sections were briefly rinsed in the same TBS/EtOH mixture, followed by a blocking step with either 3% goat or donkey serum (prepared in TBS-X) for 30min at room temperature. Sections were directly transferred to primary antibodies diluted in blocking buffer and applied overnight at 4 °C with slow agitation. The following primary antibodies were used; Aβ (3D6, a gift from Ronald Demattos, 1:1000), IBA1(1:500, 011-27991 FUJIFILM WAKO, NB100-1028, Novus Biologicals), GFAP (1:500, 53-9892-82,Invitrogen), CD31(1:250, MAB1398Z, Milipore Sigma), CD206(1:500, MR5D3, BIO-RAD), Clec7a (1:50, mabg-mdect, InvivoGen), Trem2 (1:500) and Tmem119 (1:500, E3E1O, Cell Signaling Technology). The following day, sections were washed and incubated with appropriate secondary antibodies (prepared in blocking buffer) for 2 h at room temperature. Secondary antibodies were selected based on the primary antibody host and channel of interest and used at dilutions of 1:250, 1:500, or 1:1000. Following additional washes in TBS-X and TBS, the sections were mounted onto glass slides, briefly counterstained with TrueBlack (Biotium; 23007), washed twice in MiliQ, air-dried, and coverslipped with ProLong Gold mounting medium (Thermo Fisher Scientific, P36930). Slides were dried overnight at RT and stored in the dark at 4°C until imaging.

### Imaging and quantification

For percent area analysis, slides were scanned using the THUNDER imaging system (DMi8, Leica) using 20X lens magnification. Images were further processed and quantified using Fiji software (version 2.16, ImageJ). Regions of interest (ROI) were created to select the cerebral cortex using tissue landmarks, and then a single appropriate threshold value was determined based on a preliminary analysis of at least n = 5 to consistently highlight the positive fluorescence signal. X34^+^ or Aβ^+^ CAA were manually distinguished from the parenchymal plaques based on morphology and confirmed by the presence of a positive CD31 signal. CAA was further separated into the leptomeningeal (pial) and cerebral (penetrating) vessels compartments.

For the quantification of APOE, CD206, or glial cell signals around the plaques and CAA, images were captured with 40x lens (oil) with a resolution of 524 pixels by 524 pixels and a line average of 2 using a Leica STELLARIS 5 confocal microscope and the Leica Application Suite X software (version 4.2.1.23810). 2, 3D-Z stacks, 1.5 µm step increments in the z-plane, were acquired from parenchymal plaques, pial vessel, and penetrating vessel per slice, 4 images per mouse total per region. For each region, separately, quantification of APOE, CD206, GFAP, IBA1, TMEM119, Trem2, and Clec7a was performed using a semi-automated workflow implemented in MATLAB and Imaris (v9.3.1). Surfaces were generated for the X34 signal and expanded by 15μm in the parenchymal plaques and 10μm around the pial and penetrating vessel CAA. Then, surfaces for each marker were generated, and the number of corresponding marker-positive surfaces within the defined plaque vicinity was quantified.

### Parenchyma and vessel fractionation

To quantify Aβ40/42 and APOE levels in the parenchymal and vascular compartments separately, parenchyma and vessels were fractionated using a modified version of a previously described protocol(60, 61). The procedure was scaled down for one-quarter of the mouse brain cortex tissue. Briefly, brain tissue was weighed and mechanically homogenized by repeated aspiration through a 21-gauge cannula in ice-cold buffer (HBSS supplemented with phosphatase inhibitors (PhosSTOP) and protease inhibitor cocktail), followed by centrifugation to separate the parenchymal supernatant from the vascular pellet. The supernatant, corresponding to the HBSS-soluble parenchymal fraction, was further centrifuged at high speed to separate soluble and insoluble parenchymal components. Soluble parenchymal fractions were collected from the supernatant, while insoluble parenchymal pellets were retained for further extraction. The pellet containing brain vessels was subjected to sequential buffer washes and myelin removal, followed by filtration through a 20-µm cell strainer to isolate purified vessels.

The vascular pellet was resuspended in buffer, homogenized by vortexing and pipetting, and sonicated at 20% amplitude (four 5s pulses) at room temperature using a sonicator (model FB120, Fisher Scientific). Samples were then centrifuged at 50,000g for 20 min. The resulting supernatant was collected as the soluble vascular fraction, and the pellet was processed for insoluble fractionation. For the extraction of insoluble material, parenchymal and vascular pellets were resuspended in formic acid (FA), mixed by vortexing and pipetting, and then sonicated at 20% amplitude (four 5s pulses). This was followed by centrifugation at 50,000g for 20 min. All fractions were stored transiently at −20 °C or at −80 °C for long-term storage until downstream analyses.

### Enzyme-linked immunosorbent assay (ELISA)

The levels of Aβ40, Aβ42, and APOE were measured in the HBSS- and FA-soluble fractions of the parenchymal and vascular fractions using in-house antibodies by sandwich ELISA. Briefly, the high-binding 96-well plates (half-area) were coated with the capture antibodies for either Aβ40 (anti-Aβ35-40 HJ2, in-house, 20 μg/mL), Aβ42 (anti-Aβ37-42 HJ7.4, in-house, 20 μg/mL), and APOE (HJ15.3, in-house, mouse monoclonal 5μg/mL) overnight at 4°C. The following day, the plates were washed 5 times with PBS and blocked with 2% BSA in PBS for 1h at 37 °C. Then, after another wash step in PBS, the samples and standards diluted in sample buffer (0.25% BSA + 300 mM Tris, pH 8.0 + 1x PI in PBS) were loaded in duplicate and incubated at 4°C overnight. The next day, after a series of 5 washes in PBS, the detection antibody was added and incubated at 37°C for 90 min: APOE (HJ15.7-biotinylated, in-house, 100 ng/mL) and Aβ40 and Aβ42 (anti-Aβ13-18 HJ5.1, in-house, 100 ng/mL). Plates were then washed with PBS and incubated with Strep-poly-HRP40 (1:16,000, Fitzgerald) for 90 min at RT. After additional washes, the signal was developed using Super Slow ELISA 3,3′,5,5′-tetramethylbenzidine (Sigma), and absorbance was measured at 650 nm. At least three readings were taken, and the results were plotted as the average of the three.

### Microhemorrhage analysis

Microbleeds were quantified using Perl’s Prussian blue dye to stain hemosiderin deposits as previously described, with minor modifications(20). Briefly, 5-7 equally spaced sagittal brain sections from each mouse were mounted on charged glass slides. Slices were dried overnight. First, permeabilization was performed in (0.25% Triton-X 100 in PBS) for 10 min, followed by staining for bleeds in potassium ferrocyanide (5% w/v, P3289; Sigma-Aldrich) prepared in 2% hydrochloric acid (v/v) for 30 min. The slides were then washed in water to remove excess stains. Next, the sections were stained for Aβ in 0.025% ThioS (T1892; Sigma-Aldrich), prepared in a 50:50 mixture of ethanol and TBS, for 10 min. Following a series of washes in 50% ethanol in TBS (2 × 10 min) and TBS (3 × 5 min), slides were dried and coverslipped with Fluoromount-G. All sections were imaged on a Nanzoomer 2-HT slide scanner (Hamamatsu, Bridgewater, NJ). Quantitative image analysis of microbleeds was performed using digital pathology software (Visiopharm, Broomfield, CO) with a custom-written algorithm. Investigators were blind to the treatment groups.

### Photoacoustic microscopy

Mice were anesthetized with 4% isoflurane before the craniotomy procedure and maintained under 1% isoflurane throughout the procedure. First, a longitudinal incision was made along the midline suture to expose the skull. Then, a rectangular cranial window (3 × 3 mm²) was carefully created to allow imaging of the leptomeningeal blood vessels. Mice were weaned off isoflurane and then injected with a Ketamine/Xylazine cocktail (KX, ketamine: 100 mg/kg; xylazine: 10 mg/kg). They were positioned on the stage for photoacoustic imaging, and a continuous supply of medical air was maintained. If required during imaging, mice were given a half dose of the KX cocktail. Body temperature was monitored and maintained at approximately 37 °C throughout the imaging session using a thermal probe. Mice were imaged at baseline in raster scanning mode with a laser pulse repetition rate of 6 kHz and a pulse energy of 100 nJ. Dual-wavelength (532 nm and 558 nm) scanning was applied for the measurement of blood oxygen saturation (sO_2_). Images were acquired over a 2 mm × 2 mm field of view using a step size of 0.8 µm in the X direction and 10 µm in the Y direction. Mice were then subjected to hypercapnia (10% CO_2_ in medical air for 10 min), followed by a second imaging session of the same region using the same parameters. Vascular function parameters, including blood vessel diameter, flow speed, and oxygen saturation, were quantified in response to hypercapnia using established spectral and correlation-based analyses, as previously described(44). Mice were then injected with CD31 (retro-orbitally) to stain the blood vessels, following which they were given a lethal dose of KX. The brain was topically stained for X34 for 30 minutes. After rinsing the exposed brain with PBS, the cranial window was filled with PBS, and a coverslip was mounted. A 300 µm z-stack with a step size of 12 µm was acquired using a Zeiss Examiner Z1 epifluorescence microscope equipped with a 5x air lens (0.16 NA) to capture an in-focus signal over the curved brain surface. The z-stack was deconvolved with Huygens deconvolution software as previously described, and the max intensity projection was generated into a single 2D image(62). X34% area was quantified using ImageJ within a vessel-specific ROI corresponding to photoacoustic measurements. For each vascular outcome (changes in diameter, flow speed, and oxygen saturation), the relationship with local X34 burden was assessed using linear regression. Because X34 burden did not significantly correlate with any vascular parameter and showed comparable distributions across experimental groups, functional outcomes were compared without normalization to or stratification by X34 burden.

### Statistical analysis

Data are presented as mean ± SEM unless otherwise indicated. All statistical analyses were performed using GraphPad Prism (v10.6). Data normality was assessed using the Shapiro-Wilk, D’Agostino-Pearson, and Kolmogorov-Smirnov tests. For comparisons between two groups, statistical significance was determined using a two-tailed Student’s *t* test for normally distributed data or the Mann-Whitney U test for non-normally distributed data. For comparisons between more than 2 groups, an Ordinary one-way ANOVA was used with Sidak’s multiple comparisons test. Correlations between X34 amyloid coverage and vascular hemodynamic parameters were assessed using Pearson correlation and linear regression analysis. A *P* value < 0.05 was considered statistically significant.

## List of Supplementary Materials

Fig. S1 to S7

Data file S1

## Acknowledgments

We thank R. Vassar for gifting 5XFAD (line 7031); the members in the Flow Cytometry and Fluorescence-Activated Cell Sorting Core at Washington University for sorting nuclei for sequencing; and the staff at the Genome Technology Access Center at the McDonnell Genome Institute at Washington University School of Medicine for help with genomic library construction and sequencing, NCI Cancer Center Support Grant #P30 CA91842 to the Siteman Cancer Center from the National Center for Research Resources (NCRR), a component of the National Institutes of Health (NIH), and NIH Roadmap for Medical Research; the staff at the Washington University Center for Cellular Imaging-Neuro for assistance with image analysis. We are grateful to E. Reiman, G. Serrano, and T. Beach of 1309 the Banner Sun Health Research Institute Brain and Body Donation Program of Sun City, Arizona, for 1310 the provision of human Alzheimer’s disease brain tissue. The Brain and Body Donation Program has 1311 been supported by the National Institute of Neurological Disorders and Stroke (U24 NS072026 National 1312 Brain and Tissue Resource for Parkinson’s Disease and Related Disorders), the National Institute on 1313 Aging (P30 AG19610 Arizona Alzheimer’s Disease Core Center), the Arizona Department of Health 1314 Services (contract 211002, Arizona Alzheimer’s Research Center), the Arizona Biomedical Research 1315 Commission (contracts 4001, 0011, 05-901 and 1001 to the Arizona Parkinson’s Disease Consortium) 1316 and the Michael J. Fox Foundation for Parkinson’s Research.

## Author contributions

Conceptualization: FS, DMH, JDU

Methodology: FS, YC, IVK, BL, YW, LCDS, MEB, KAT, SP, XB, AN, BHZ, KH

Investigation: FS, YC, YW, KAT, MEB

Visualization: FS, DMH, JDU

Funding acquisition: DMH, JK, GJR, FS

Project administration: FS

Supervision: DMH, JDU, GJR, SH, JK

Writing – original draft: FS

Writing – review & editing: FS, DMH, JDU, YC, IVK, BL, YW, LCDS, MEB, KAT, SP, XB, AN, BHZ, GJR, SH, JK, KH

## Data and materials availability

All relevant data associated with this study are within the manuscript and its Supplementary Materials files.

## Supplementary figures

**S1.**
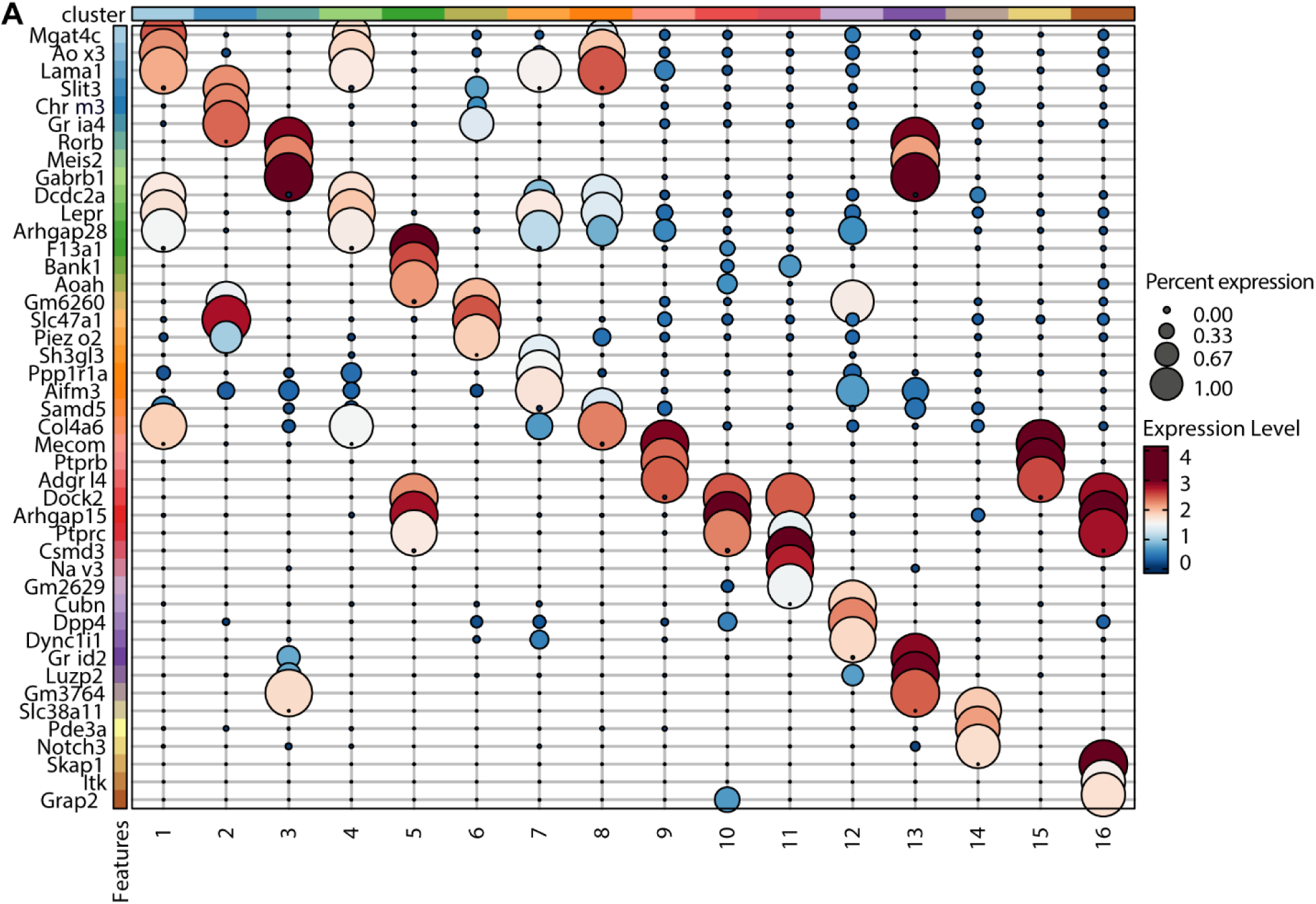
Marker gene expression across cell clusters. (A) Dot plot displaying the expression of selected marker genes used to annotate cell clusters identified in the dataset. Columns correspond to clusters, and rows correspond to genes. Dot size indicates the fraction of cells within each cluster expressing the gene, and dot color reflects the scaled average expression level.

**S2.**
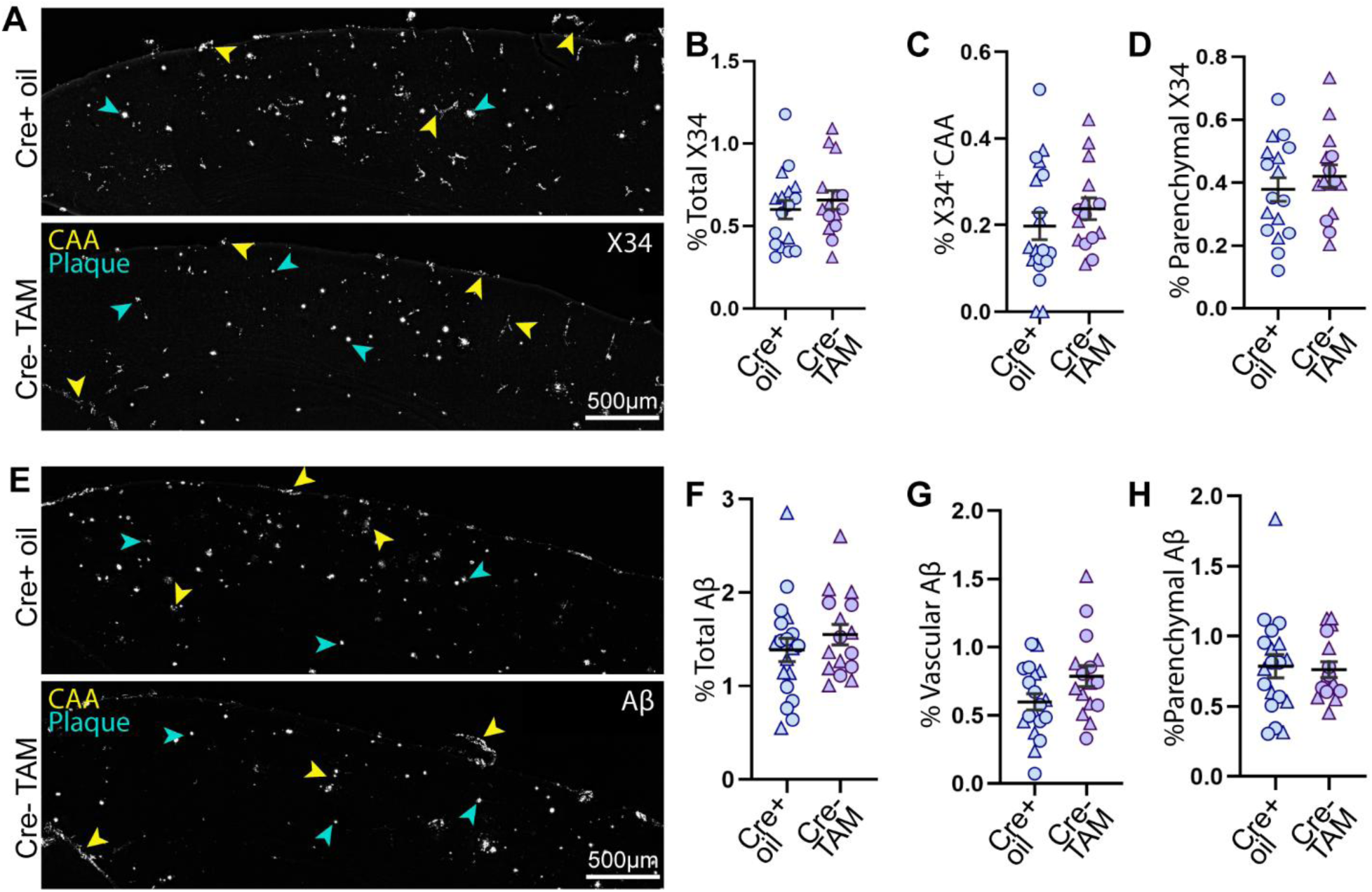
Cre expression does not affect amyloid burden or CAA. (A) Representative fluorescence images of cortices stained for fibrillar Aβ (X34). (B-D) Quantification of the percent area for total X34 (B), CAA (C), and parenchymal plaques (D). (E) Representative images stained for Aβ immunoreactivity in the cortex. (F-H) Quantification of the percent area for total Aβ (F), CAA (G), and parenchymal plaques (H). Yellow arrows indicate CAA, and teal arrows indicate parenchymal plaques. n=15-17 each group. n reflects the number of animals included per analysis after predefined exclusion criteria. All data expressed as means ± SEM. Student’s t-test with Welch’s correction.

**S3.**
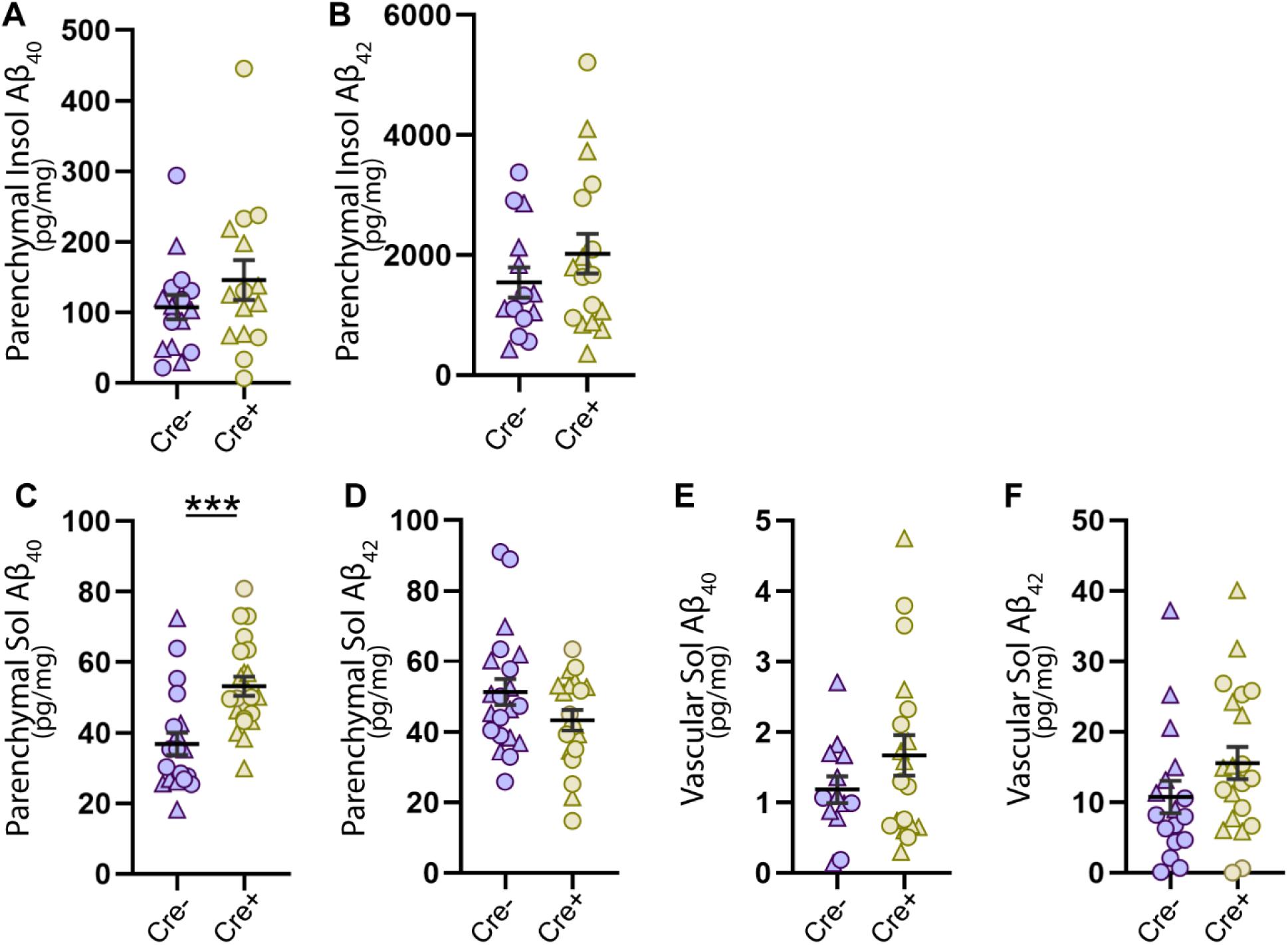
HBSS- and FA-soluble Aβ40 and Aβ42 levels in vascular and parenchymal fractions. (A, B) Aβ40 (A) and Aβ42 (B) levels quantified by ELISA in the formic acid (FA)-soluble parenchymal fraction. (C, D) Aβ40 (C) and Aβ42 (D) levels quantified by ELISA in the HBSS-soluble parenchymal fraction. (E, F) Aβ40 (E) and Aβ42 (F) levels quantified by ELISA in the HBSS-soluble vascular fraction. n= 17-18 (Cre⁺) and 15-20 (Cre⁻). n reflects the number of animals included per analysis after predefined exclusion criteria. All data expressed as means ± SEM. Student’s t-test with Welch’s correction, \*\*\**P* < 0.001.

**S4.**
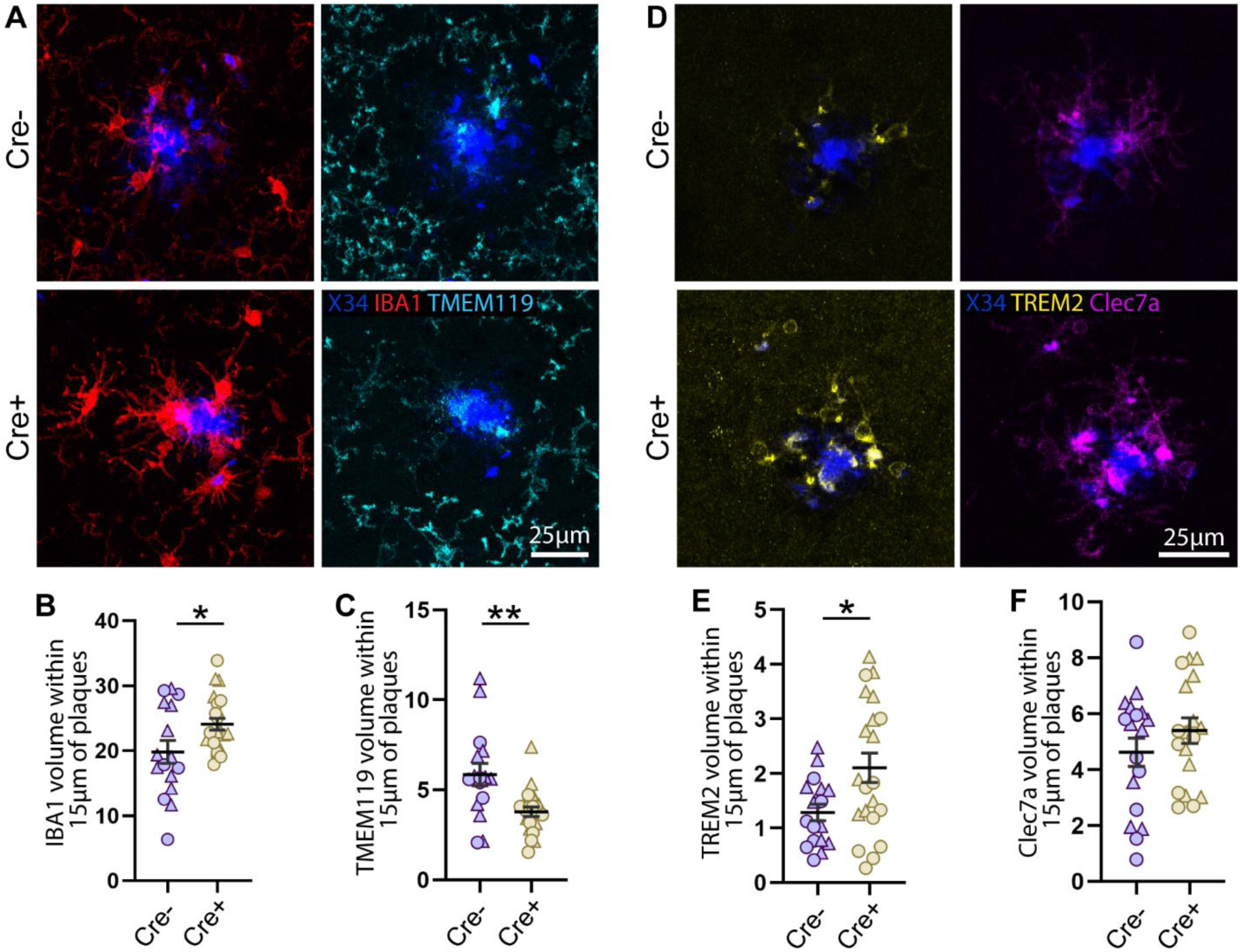
APOE4 depletion in VAFs increases microglial reactivity around parenchymal plaques. (A) Representative immunofluorescence images of fibrillar Aβ (X34) in parenchymal plaques and microglial markers IBA1 and TMEM119. (B, C) Quantification of % IBA1 (B), and TMEM119 (C) volume within a 15μm radius of parenchymal plaques shown in (A). (D) Representative immunofluorescence images of fibrillar Aβ (X34) in parenchymal plaques and microglial markers TREM2 and Clec7a. (E, F) Quantification of % TREM2 (E) and Clec7a (F) volume within a 15μm radius of parenchymal plaques shown in (D). n = 19-21(Cre⁺) and 16-18 (Cre⁻). n reflects the number of animals included per analysis after predefined exclusion criteria. All data expressed as means ± SEM. Student’s t-test with Welch’s correction, \**P* < 0.05, \*\**P* < 0.01.

**S5.**
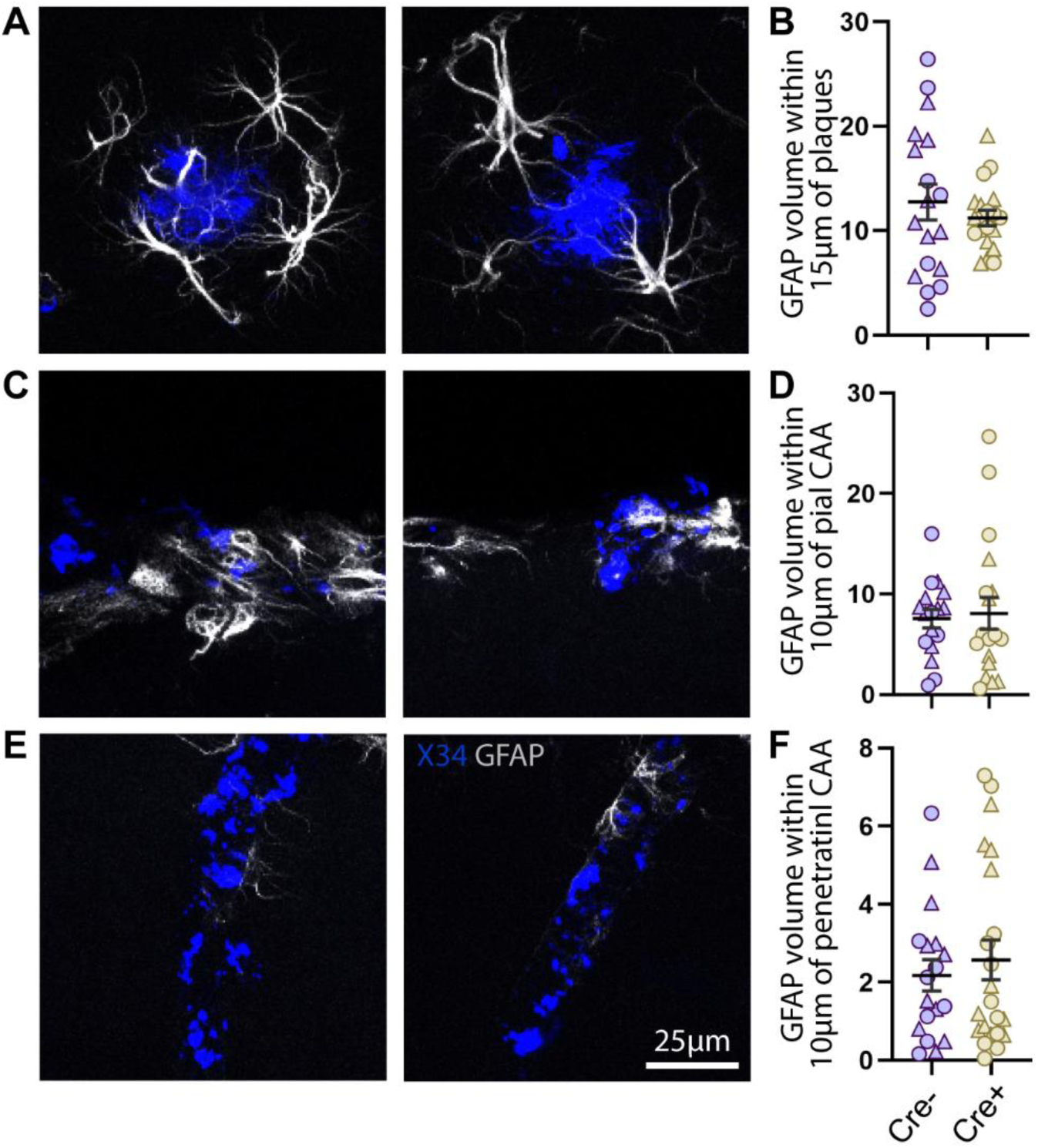
APOE4 depletion in VAFs does not affect GFAP expression around CAA and parenchymal plaques. (A) Representative immunofluorescence images of fibrillar Aβ (X34) in parenchymal plaques and astrocyte (GFAP). (B) Quantification of % GFAP volume within a 15μm radius of parenchymal plaques shown in (A). (C) Representative immunofluorescence images of fibrillar Aβ (X34) in pial CAA and astrocyte (GFAP). (D) Quantification of % GFAP volume within a 10μm radius of pial CAA shown in (C). (E) Representative immunofluorescence images of fibrillar Aβ (X34) in penetrating CAA and astrocyte (GFAP). (F) Quantification of % GFAP volume within a 10μm radius of penetrating CAA shown in (F). n = 19-22(Cre⁺) and 17-18 (Cre⁻). n reflects the number of animals included per analysis after predefined exclusion criteria. All data expressed as means ± SEM. Student’s t-test with Welch’s correction.

**S6.**
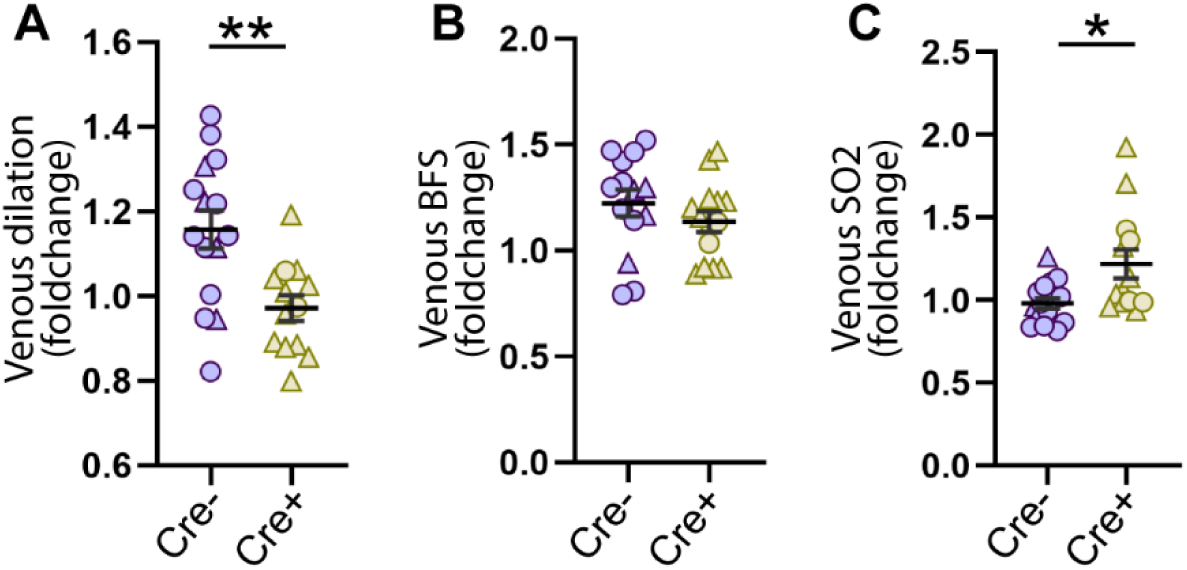
APOE4 depletion in VAFs altered pial vein function in Cre⁺ mice. (A-C) Quantification of fold-change in venous dilation (A), venous blood flow speed (BFS) (B), and venous oxygen saturation (sO_2_) (C) upon hypercapnia relative to baseline in pial veins. n = 13-14 vessels from 7 mice (Cre^+^) and 14-16 vessels from 8 mice (Cre^-^). All data expressed as means ± SEM. Student’s t-test with Welch’s correction.

**S7.**
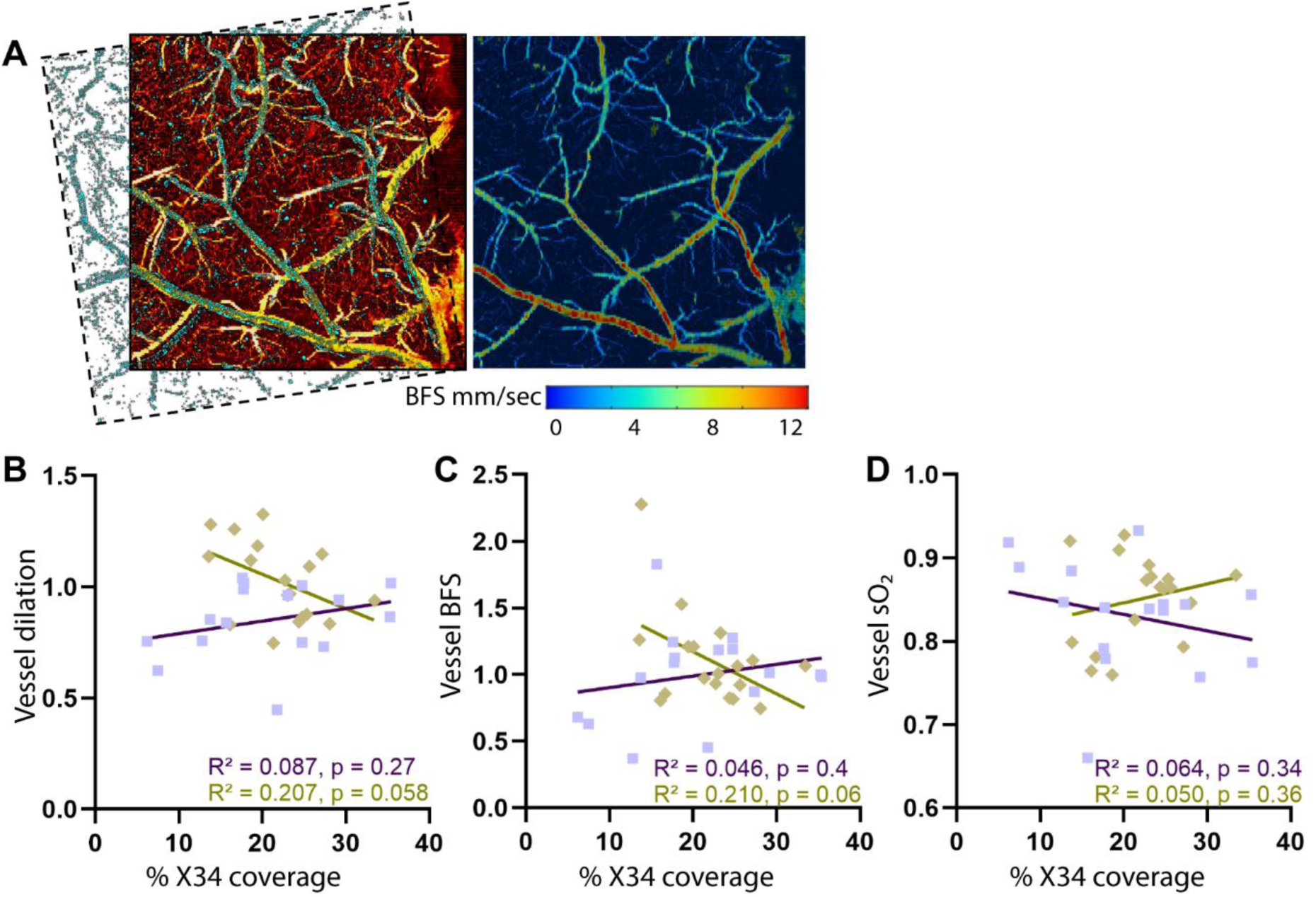
X34 amyloid burden does not correlate with vascular hemodynamic changes. (A) Photoacoustic image co-registered with X34-stained widefield images. Correlation analysis of X34 coverage with vessel dilation (B), blood-flow speed (BFS) (C), and oxygen saturation (sO₂) (D).

